# Simulation of aerosol dispersion during medical examinations

**DOI:** 10.1101/2021.11.22.469529

**Authors:** Sebastian Falk, Sarina K Mueller, Stefan Kniesburges, Michael Döllinger

## Abstract

The main route of transmission of the SARS-CoV2 virus has been shown to be airborne. The objective of this study is to analyze the aerosol dispersion and potential exposure to medical staff within a typical medical examination room during classical airway procedures. The multiphase simulation of the aerosol particles in the airflow is based on a Lagrangian-Eulerian approach. All simulation cases with surgical mask show partially but significantly reduced maximum dispersion distances of the aerosol particles compared to the cases without surgical mask. The simulations have shown that medical examiner are exposed to large amount of aerosol particles, especially during procedures such as laryngoscopy where the examiner’s head is directly in front of the the patient’s face. However, exposure can be drastically reduced if the patient wears a mask which is possible for the most of the procedures studied, such as otoscopy, sonography, or anamnesis.

## 1 Introduction

Coronavirus disease 2019 (COVID-19) is an acute respiratory illness caused by the SARS-CoV-2 (Severe Acute Respiratory Syndrome Coronavirus 2) coronavirus. The socioeconomic impact cannot be compared to any other pandemic before [1]. SARS-CoV-2 results in high variability of symptoms. While some patients have mild symptoms of upper respiratory tract infection, others suffer severe sequelae resembling acute respiratory distress syndrome (ARDS) [2–4].

The main route of transmission of the SARS-CoV2 virus has been shown to be airborne by hoovering aerosol particles. To date, at least 115,000 healthcare workers have died from SARS-CoV2 [5]. Owing to the high exposure of aerosol particles during clinical airway procedures [6, 7]. Thus, for their own safety, but also for the maintenance of healthcare and the safety of the rest of the population, it is of great importance to minimize the exposure of clinical staff. Several recommendations to ensure maximum safety of medical personnel during airway management procedures are developed and updated regularly [6–11]. This includes limiting some procedures to the minimum necessary, including bronchoscopy or tracheal cannula change in SARS-CoV2-positive patients.

Looking at the receiver, inhalable aerosol particles must contain particles or liquid droplets smaller than ≈ 20 *μm*. However, only particles with a diameter < 5 *μm* (fine particles) can effectively penetrate the lower parts of the respiratory tract, e.g., the small bronchi and alveoli [12]. There are different definitions of airborne transmission of pathogens by liquid aerosol particles. Some authors use the terms droplets for objects larger than 20 *μm* and droplet nuclei for objects smaller than 5*μm* [4, 8, 13, 14]. Other authors distinguish between liquid particles suspended in the air that remain in the air for a short time due to ballistic trajectories (droplets) and particles that float and remain in the air for a longer time (aerosol particles). In this study, based on previous results of our group (clinical characterization), we will use the second definition based on ballistic and hovering behavior of respiratory particles [15]. Filtering mechanisms in the upper airways effectively seems to remove the majority of particles larger than 5 *μm* from the air stream already in the nose/mouth/throat [12]. After passing through the upper airway filters, aerosol particles enter the bronchial tree more slowly. Particles that do not settle during inhalation may also be retained in the airways during exhalation [12]. In this context, it is important to emphasize that aerosol particles from the bronchial tree/alveoli can increase their size in the airways and be delivered as droplets on exhalation. Ultrafine particles (< 100 *nm*) have a high probability of deposition to the bronchial tree and alveoli. However, they are not considered because coronavirus, an enveloped virus with a size of 120 – 160 *nm*, is larger than ultrafine aerosol particles and were found to be clinically insignificant [4, 12, 16]. Exhaled aerosol particles that are between 0.1 and 0.5 *μm* in size may potentially contain at least one virostatic agent [4, 12, 16].

A prerequisite for infection is that the virus remains infectious and replicable in these small particles [14]. Previous studies have shown that the SARS-CoV2 virus can remain airborne for several hours [17]. However, the airborne residence time depends on size and trajectory, among other factors. The aerosol production rate varies greatly from person to person. While breathing rate has been shown to have no effect on aerosol concentration, inhalation of surfactant or isotonic NaCl results in significant changes. Inhalation of surfactant increased the aerosol concentration by 300 %. Inhalation of isotonic NaCl, on the other hand, was able to reduce particle exhalation by < 70 % [18]. A study by [19] also indicated that patients with respiratory infections produce more aerosol particles (especially at 1 *μm*) than healthy individuals [11, 19, 20]. While gender was not a major influence in previous studies, the product of age and body mass index (BMI) turned out to be a good predictor of aerosol production with a critical threshold for increased aerosol production of 650 and larger [21].

Numerous studies on the fluid dynamics of droplets and aerosol particles have been conducted in recent years, particularly in the context of the COVID-19 pandemic.

Experimental studies focusing on the behavior of cough flow at greater distances from the mouth [22] or on the droplets and aerosol particles during normal speech [23] were done.

Numerical studies on the effect of wind speed on social distancing [24], of high flow oxygen therapy [25], and of droplet behavior at various relative humidities were performed [13]. Several other studies (e.g., [26–29]) and international online conferences [30, 31] demonstrate the usefulness and importance of computational fluid dynamic (CFD) simulations for the current COVID-19 pandemic.

An experimental study using particle image velocimetry and hotwire anemometry was conducted in combination with CFD simulations [22]. In situ measurements and numerical simulations were combined and focused on exhaled aerosol particles under normal respiratory behavior and their transport in an elevator and in small classrooms [32]

Investigations of droplet or aerosol dispersion during normal speech [23, 33], coughing [33, 34], or breathing [33] were done.

The effects of using a type 1 surgical mask over a patient’s face on particle behavior [25] and that masks were able to protect individuals in the environment during breathing, speaking, singing, coughing, and sneezing [33] were studied.

However, these data were not collected under real conditions during airway management procedures with patients. The bulk of the described studies did not deeply investigate airway management procedures.

Thus, it is the aim of this study to analyze the aerosol dispersion and potential exposure to medical staff within a typical medical examination room during classical airway procedures. Detailed knowledge of the size, velocity, trajectories, and flow patterns of various upper and lower airway management procedures is essential for developing effective strategies to reduce the exposure of aerosol particles for medical personnel. Currently, there is no study and no quantitative real-life data.

## 2 Methods

The simulations were performed with Simcenter STAR-CCM+ 2020.2 CFD solver (Siemens, PLM Software, Plano, TX, USA) based on a Lagrangian Euler approach to model the multiphase flow of the aerosol dispersion [35]. The model uses a combination of an Eulerian segregated solver for the continuous phase in conjunction with a Lagrangian multiphase model for the dispersed phase [35]. The Eulerian-Lagrangian approach exhibited good agreement to experimental data as shown in [36].

Furthermore, the Lagrangian-Eulerian model additionally considers heat and mass transfer phenomena [37]. Coupling between the two phases occurs by mass, momentum, and energy transfer [37]. In the study, one-way coupling was used, where only the continuous phase affects the dispersed phase as exclusively small particles were examined which follow the convectional flow in a slip-free environment having a Stokes number (St) much lower than one (St << 1) [38].

For modeling turbulence, the unsteady Reynolds-averaged Navier-Stokes model (unsteady RANS) is used with the well-established and widely used SST (Menter) K-Omega model [39].

A constant density is assumed for the continuous and dispersed phases. The airflow in the continuous phase has a density of 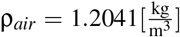, and a dynamic viscosity of 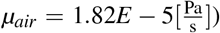, and the aerosol particles of the dispersed phase are defined as liquid particles with the properties of water/slime with a density of 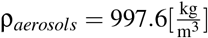[40].

In order to track the aerosol behavior over a sufficiently long and appropriate time period in the numerical domain, a physical time of 60 s is simulated. A second-order upwind scheme is used to discretize the convective and diffusive terms of the Navier-Stokes equations [41]. The implicit method is used for this transient simulation. Time step sizes of 10^−3^ s for the physical time of 0 s < *t* ≤ 10 s and of 10^−2^ s for 10 s < *t* ≤ 60 s are used [25, 42–44]. An algebraic multi-grid method with a Gauss-Seidel relaxation scheme is used to solve the final linear system of partial differential equations.

### 2.1 Simulation cases

The eight cases of the five paradigm (*anamnesis*, *laryngoscopy*, *otoscopy*, *rhinoscopy*, and *sonography*) in combination with the three inlet profiles (speaking, breathing, and coughing), see table 1, with and without a surgical mask are simulated for a physical time of 60 s. The paradigm mainly define the position of the medical examiner in relation to the patient. Overall 16 simulation cases were executed.

**Table 1.** The paradigm in combination with their inlet profiles, which were performed in this study. A total of 16 simulation cases for eight paradigm (with and without surgical mask) were simulated.

| Case | Paradigm | Position: Patient and Examiner | Inlet profile |
| --- | --- | --- | --- |
| 1 | *Anamnesis* | sitting / standing in distance | Speaking |
| 2 | *Laryngoscopy* | sitting / standing in front | Breathing |
| 3 | *Laryngoscopy* | sitting / standing in front | Coughing |
| 4 | *Otoscopy* | sitting / standing besides | Breathing |
| 5 | *Otoscopy* | sitting / standing besides | Speaking |
| 6 | *Rhinology* | sitting / standing in front | Sneezing |
| 7 | *Sonography* | lying / sitting besides | Breathing |
| 8 | *Sonography* | lying / sitting besides | Speaking |

From the simulations, the following parameters are calculated for different aerosol sizes: Dispersion of aerosol particles near the investigator’s head and the extent in all three spatial directions. The values are compared descriptively between the simulation cases.

### 2.2 Numerical domain

The numerical domain of the examination room consists of a rectangular room with a floor plan of 3.0 x 4.5 m and a ceiling height of 2.5 m, see Fig. 1. The patient sits on a treatment chair or lies on a treatment couch depending on the respective paradigm, see table 1. The medical examiner is placed in front or at the side of the patient, see Fig. 1.

**Fig. 1.**
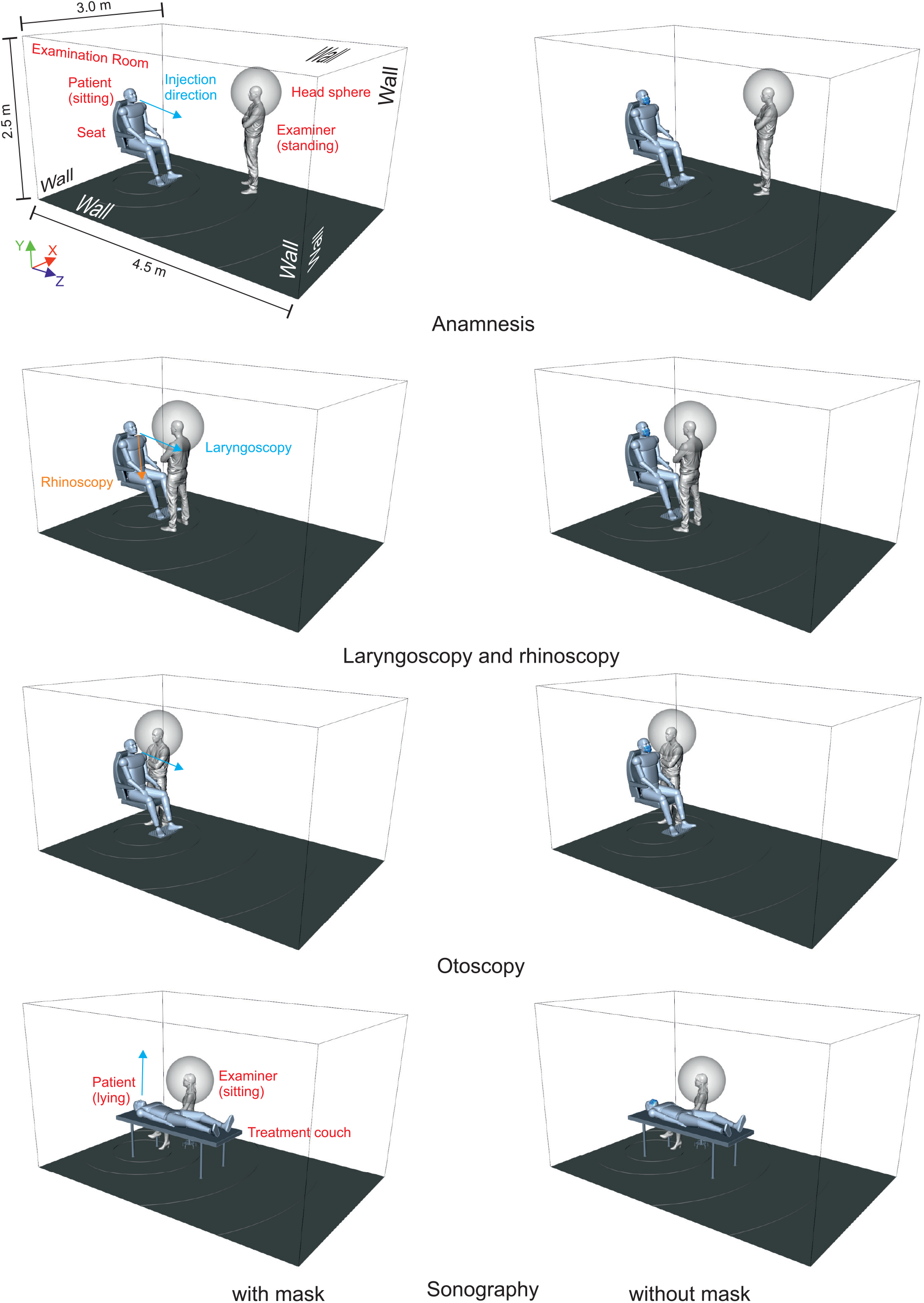
General geometry of the numerical examination room of all simulation cases and the positions of the patient and the examiner within it. The head spheres are positioned around the examiners’ heads with a diameter of 0.8 m and their surfaces track the incoming aerosol particles.

### 2.3 Boundary conditions

All surfaces including walls, persons and furniture are defined as non-moving impermeable walls, see Fig. 1. The surgical masks were considered as impermeable rebound walls. The patient’s mouth is defined as mass (speaking and breathing) or velocity (coughing) inlet for the airflow of the continuous phase. For the *rhinoscopy* the inlet is the nose (sneezing).

There is no initial velocity of the environmental air in the examination room. However, the flow fields of the continuous phase are initialized by simulating 60 s physical time using the respective profile inlets. The flow velocity during the experimental procedures ranges in the low Reynolds number (Re < 1000) region with the kinematic viscosity of air 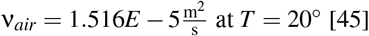.

#### Profiles of the inlet boundaries (continuous phase)

The realistic coughing velocity profile of Gupta et al. [46] is scaled to the maximum total velocity of 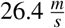 taken from the experiments of Müller et al. [15], see Fig. 2. This cough profile is then used for the paradigm with coughing and sneezing for *rhinoscopy*. The breath volume flow profile, defined as mass flow, is taken from Tabuenca Javier Garcia [47], see Fig. 2. Continuous speech was defined by a mass flow of 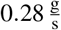 [48].

**Fig. 2.**
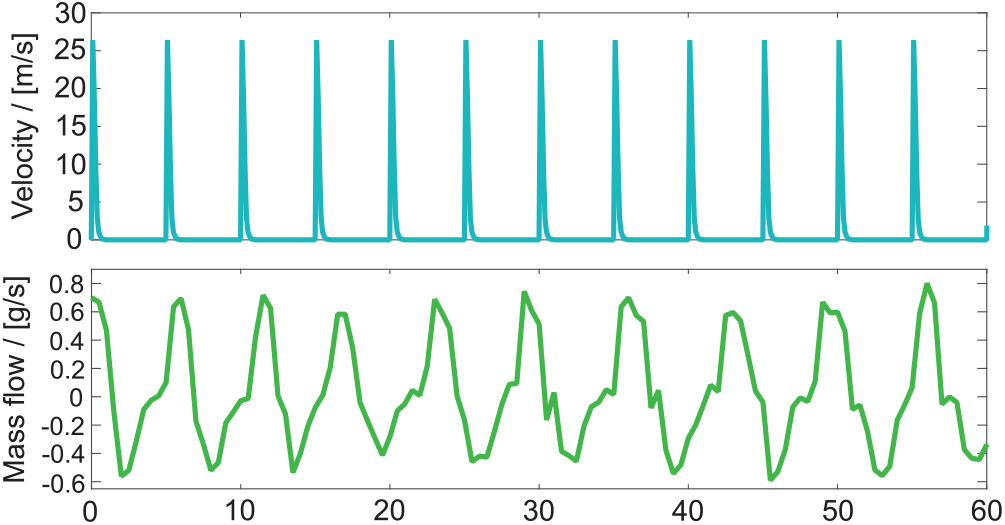
Profiles of coughing/sneezing (upper part) and breathing (lower part) for 60 s.

For the mesh independence study an polyhedral grid with an adaptive meshing strategy was used, see Fig. 3. Similar to Pendar and Páscoa [34], an inlet velocity at the patient’s mouth (inlet boundary) was specified time-dependently on the basis of the cough profile. Five different meshes (MB – M4) with an increasing spatial resolution (1.99 – 10.30 million elements) were created. Mesh M2 with a tolerable deviation of the velocity at measuring point P1 at a distance of 0.1 m from the inlet and a calculation time kept within limits was selected, see table 2.

**Fig. 3.**
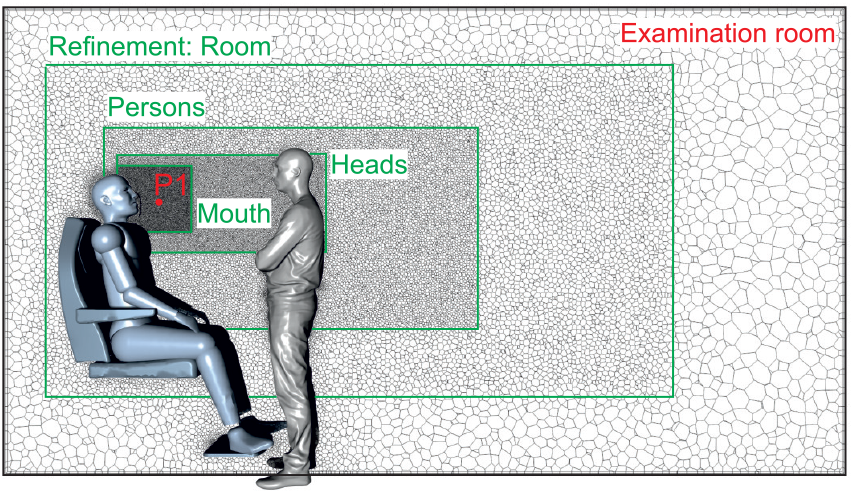
Grid within the numerical examination room with the four refinements steps and the velocity measuring point P1.

**Table 2.**
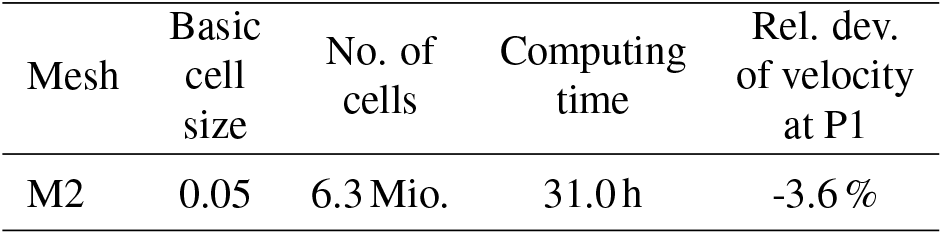
Properties of mesh M2 of the grid independence study. The velocity at measuring point P1 is given as relative deviation to mesh MB.

### 2.4 Particle distribution and injection (dispersed phase)

Experimental measurements are used as input for the injected aerosol particles in the simulation cases. The particle distributions during breathing, coughing, and speaking are based on experimental measurements with an Optical Particle Sizer (OPS) (Model 3330, TSI Incorporated, Shoreview, Minnesota, USA). The OPS divides the measuring range into so-called size buckets.

Based on the measurements, Fig. 4 shows the relative number of aerosol particles distribution for breathing, speaking and coughing for the OPS buckets. For each bucket, the mean value of the specific range and the percentage of aerosol particles are given. For breathing, the OPS detected aerosol particles only in the first 5 buckets, for speaking in the first 10 buckets, and for coughing in 12 buckets. These size distributions were similarly reported in the literature [49].

**Fig. 4.**
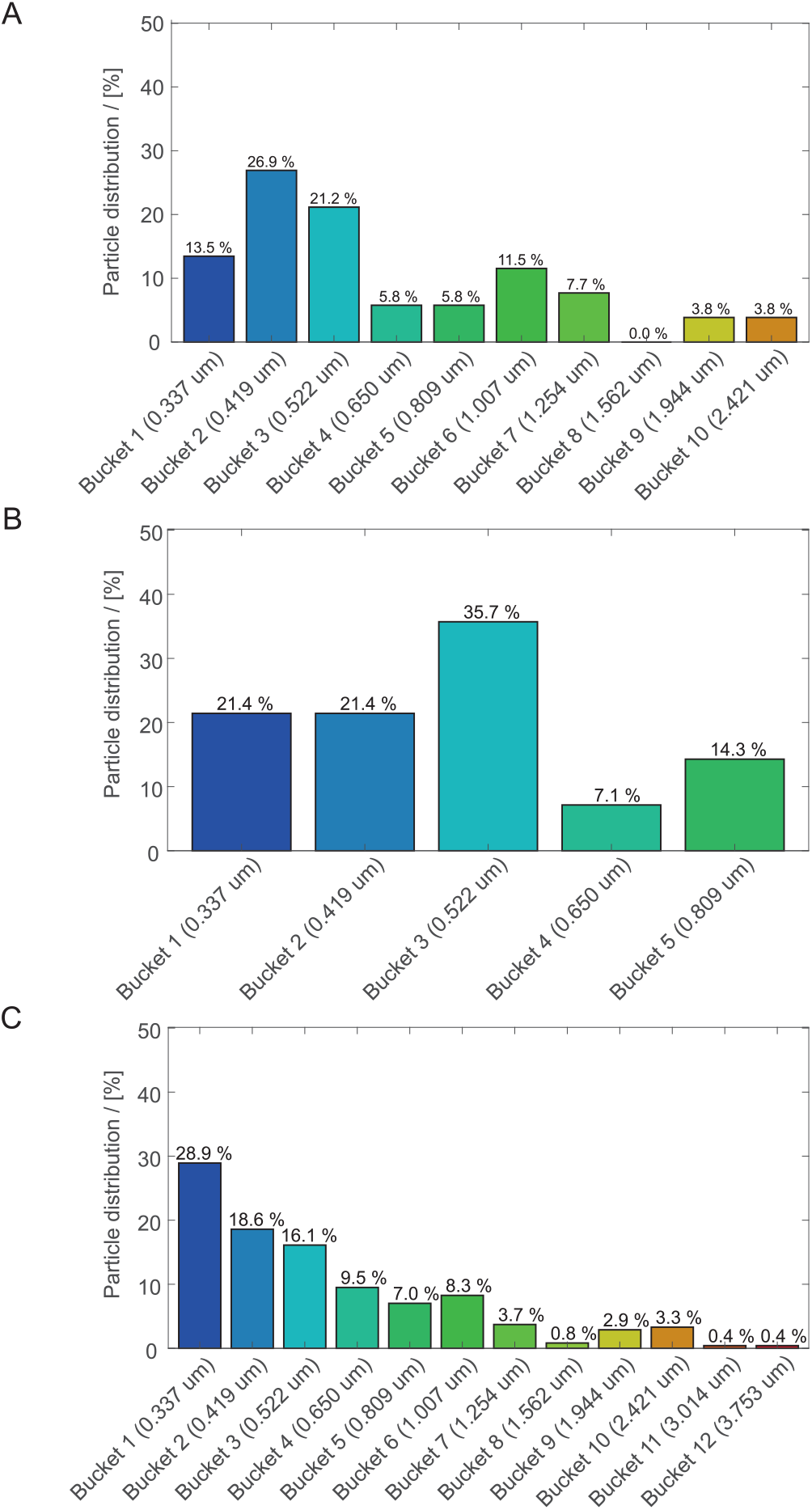
Percentage particle distribution of (A) breathing, (B) speaking, and (C) coughing among the buckets of the OPS.

At the inlet boundary, an injector is placed which feeds the aerosol particles with size distribution from Fig. 4 into the simulation region.

The inject axes, see Fig. 1, are along:

- the z-axis for *anamnesis*, *laryngoscopy*, and *otoscopy*,
- the y-axis in positive direction for *sonography*,
- the y-axis in negative direction for *rhinoscopy*.

The injector forms a cone-shaped spreading characteristic with a inclination angle of 60°. The aerosol particles are injected with an initial velocity of 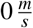 and subsequently captured by the inlet flow which is not decelerated by the particles.

## 3 Results

### 3.1 Dispersion of aerosol particles on the examiners head sphere

To count the particles that enter the near field of the examiner’s head, we created so-called head spheres around the examiner’s head. Figure 1 shows for each simulation configuration the head spheres of diameter 0.8 m around the examiners’ heads. Aerosol particles that cross the head spheres were tracked and counted.

Table 3 shows that the head spheres of the cases *laryngoscopy* with breathing and coughing, each without surgical mask, registered an enormous amount of aerosol particles with 55.0 % and 64.0 % of the total number of aerosol particles emitted by the patient. The head sphere of *otoscopy* breathing with surgical mask counted a relatively small number of aerosol particles of about 0.03 %.

**Table 3.** Percentage of the total number of particles fed into the numerical domain of the 16 simulation cases that were detected by the examiners’ head spheres.

| Case | Paradigm/Profile | with surgical mask | without surgical mask |
| --- | --- | --- | --- |
| 1 | Anamnesis/Speaking | 0.0 % | 0.0 % |
| 2 | Laryngoscopy/Breathing | 0.0 % | 55.0 % |
| 3 | Laryngoscopy/Coughing | 0.0 % | 64.0 % |
| 4 | Otoscopy/Breathing | 0.03 % | 0.0 % |
| 5 | Otoscopy/Speaking | 0.0 % | 0.0 % |
| 6 | Rhinology/Sneezing | 0.0 % | 0.0 % |
| 7 | Sonography/Breathing | 0.0 % | 0.0 % |
| 8 | Sonography/Speaking | 0.0 % | 0.0 % |

Figure 5 shows the size distribution for the three simulation cases that registered aerosol particles at the examiners’ head spheres. The largest percentage of the tracked aerosol particles occurs with low diameter particles.

**Fig. 5.**
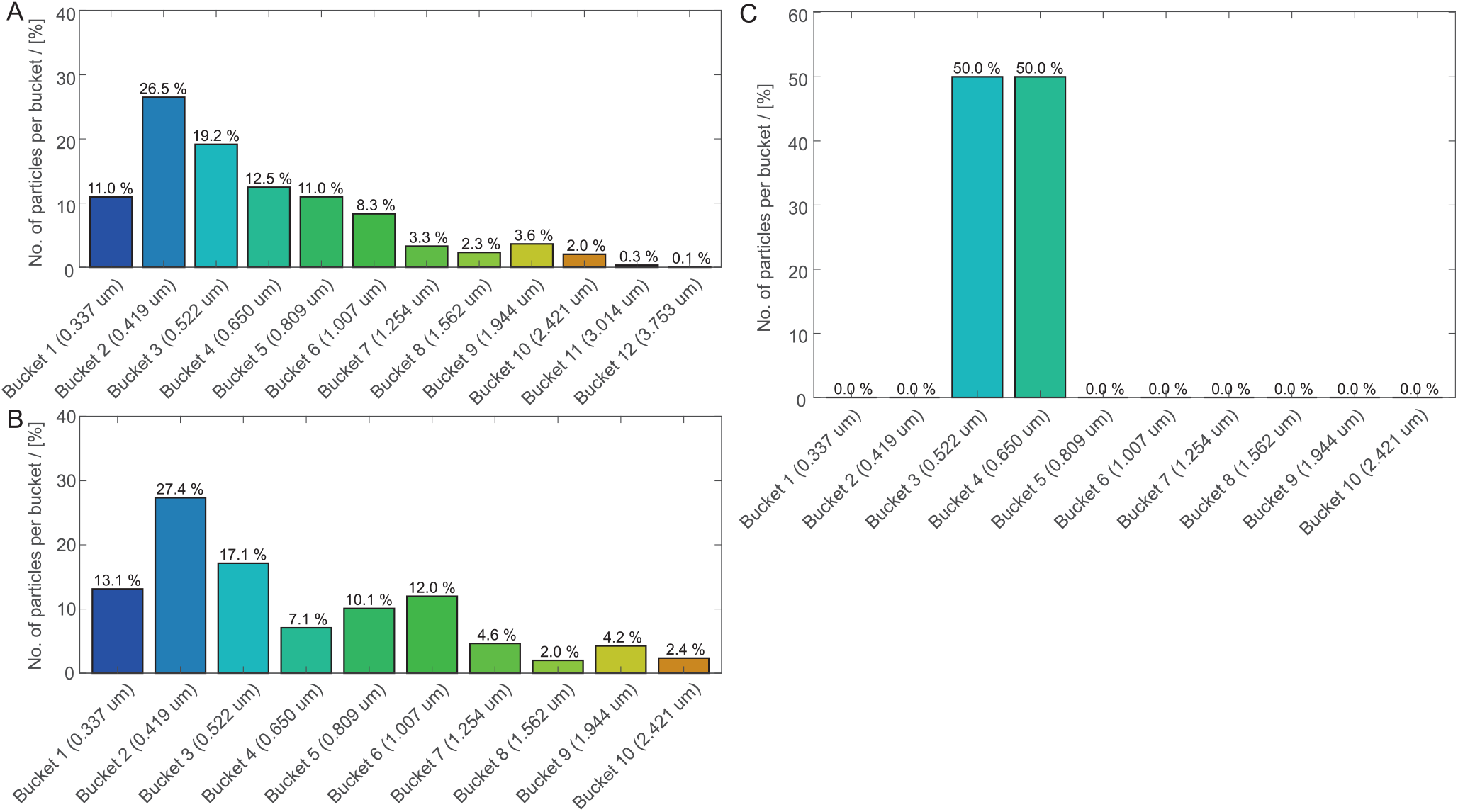
Distribution of aerosol particles entering the examiner’s head sphere by size for (A) *laryngoscopy*/without surgical mask/coughing, (B) *laryngoscopy*/without surgical mask/breathing, and (C) *otoscopy*/with surgical mask/breathing.

### 3.2 Dispersion of aerosol particles in the examination room

Figure 6 shows the maximum dispersion in all three spatial directions of the aerosol particles in the examination room. Additionally, see supplement videos for the dispersion of the aerosol particles in the examination room over time. All simulation cases with surgical mask show partially but significantly reduced maximum dispersion distances of the aerosol particles compared to the cases without surgical mask.

**Fig. 6.**
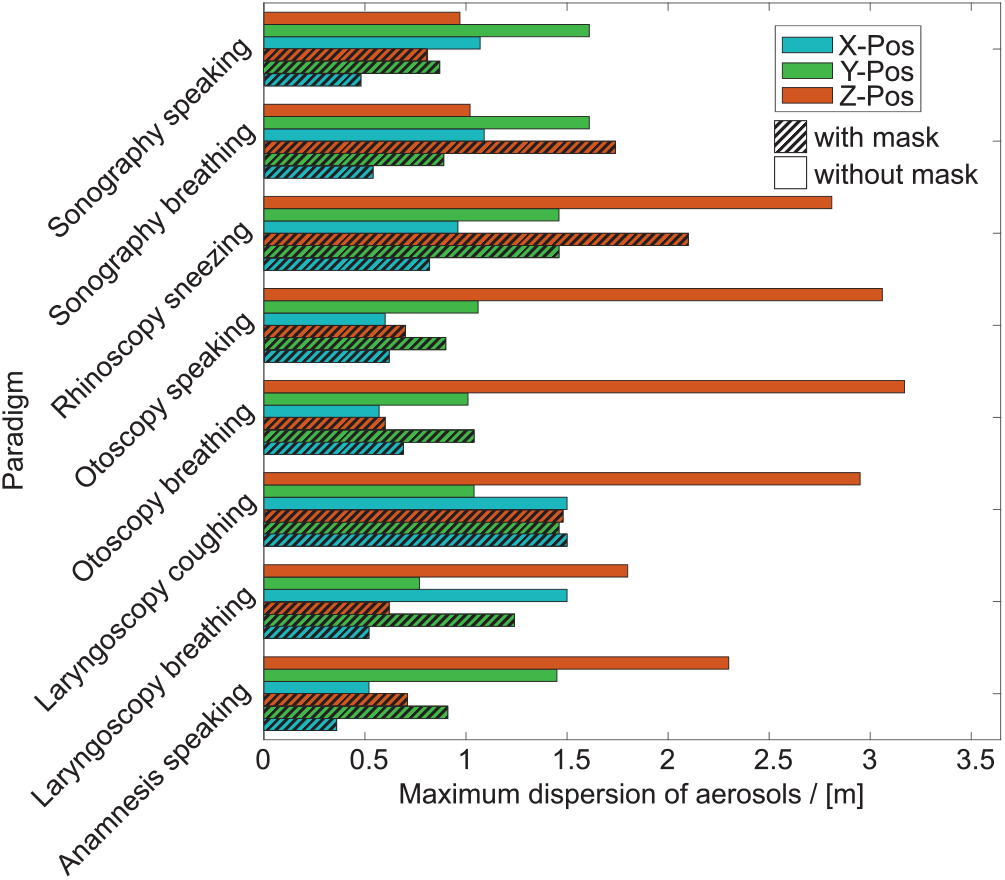
Maximum dispersion (in x-, y-, and z-direction) of the aerosol particles in the numerical domain for each paradigm with and without surgical mask.

The main direction without mask is the z-direction which is the main direction of ejection except of the case *sonography* where the patient lies on a couch and ejects upward in y-direction. Using a mask deflects the aerosol particles in orthogonal directions to the main ejection direction. For *otoscopy*, these are the positive and negativ x- and y-directions. In *sonography*, the particles are deflected in positive and negative z-instead of y-directions by the mask. For *rhinoscopy* sneezing the main ejection is forward-downward directed as shown in supplement video V11.

Figure 7 shows the maximum travel distance (length of the vector from the injector to the material particle) reached by the aerosol particles within a physical time of 60 s in the numerical domain. It can be clearly seen that the maximum travel distance of aerosol particles with larger size (≥ 2.241 *μm*) decreases with an increasing particle diameter. Furthermore, all simulation cases with surgical mask show a significantly reduced maximum travel distance of the aerosol particles among all buckets.

**Fig. 7.**
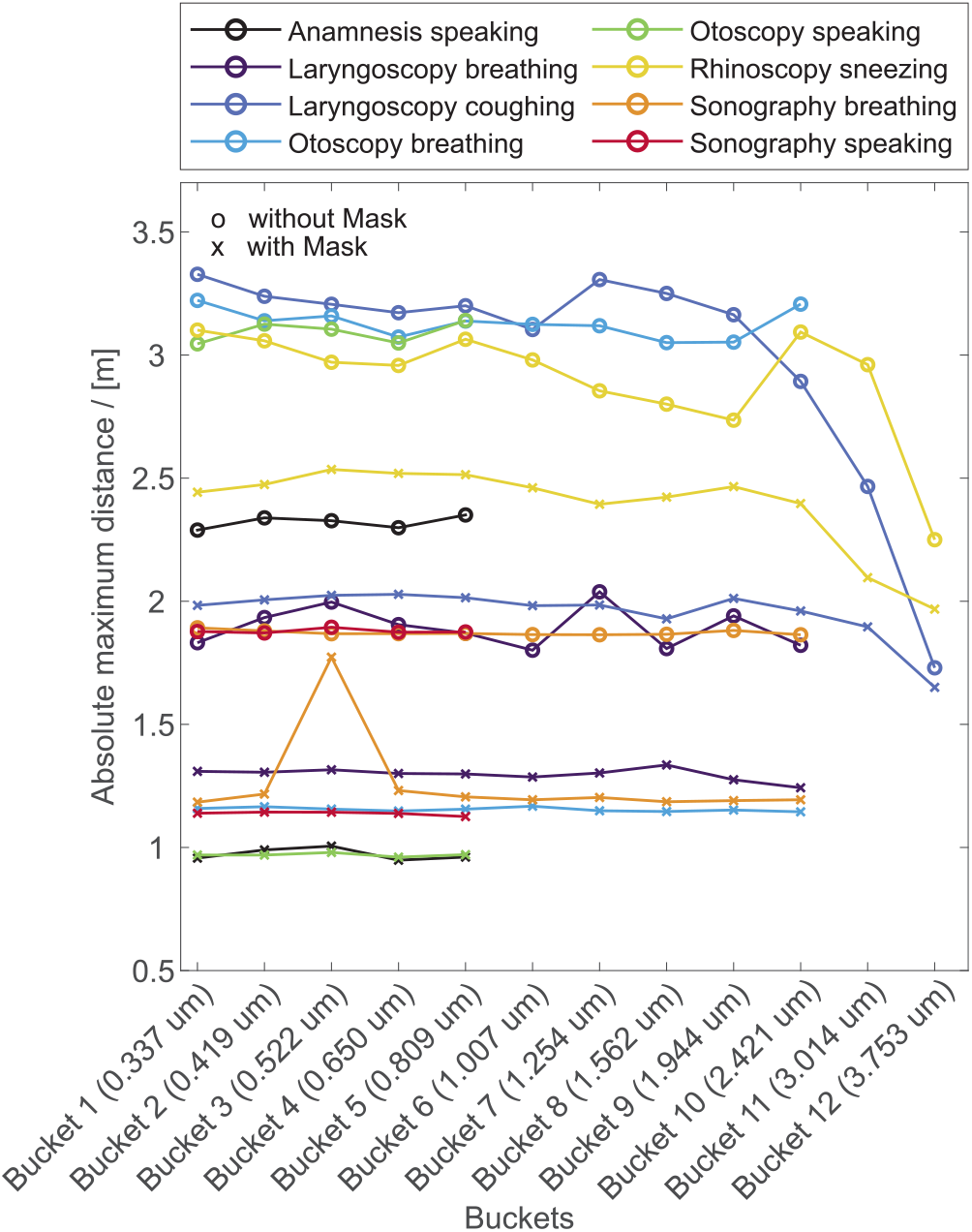
Absolute maximum distance the aerosol particles travel in the numerical domain.

## 4 Discussions

The large amount of aerosol particles delivered to the near field of the examiner’s head in the cases *laryngoscopy* with breathing and coughing is a consequence of the examiner’s position in a short distance in front of the patient’s face. The more the examiner is positioned within the patient’s exhalation flow region, the more aerosol particles will reach the near field of the examiner especially for coughing. The reason for the detected aerosol particles during *otoscopy* breathing with surgical mask is, that the exhaling flow is redirected by the surgical mask in lateral direction, so that a small amount of aerosol particles are carried by the flow and were able to enter the head sphere of the examiner, see Fig. 1. This deflection of aerosol particles by different types of masks was multiply observed in experiments [50–52].

The main direction without mask is the exhalation direction for all paradigms and profiles. After a physical time of 60 s, no significant difference in the maximum aerosol dispersion between coughing, breathing and speaking were observed. The maximum travel distance of the aerosol particles in the exhalation direction is significantly reduced by wearing a surgical mask. In addition, larger aerosol particles (≥ 2.241 *μm*) appear to travel less distance compared to smaller aerosol particles.

## 5 Conclusions

The simulations showed that medical examiner are exposed to large amount of aerosol particles especially for interventions as *laryngoscopy* where the examiner’s head is immediately in front of the the patient’s face. However, the exposition can be drastically reduced if the patient wears a mask which is in principle possible for the most investigated interventions as *otoscopy*, *sonography* or *anamnesis*. Even for *rhinoscopy*, the patients should be able to wear a mask. Unfortunately, wearing a mask by the patient is hardly an option for *laryngoscopy* as the examiner has to position the laryngoscope safely in the oral cavity without irritating the pharyngeal region, which would force coughing or even vomiting. In fact, these two effects often occur during *laryngoscopy*.

As a consequence of these findings, the examiner should be outside the patient’s exhaled air stream, regardless of whether the patient is coughing, speaking, or breathing. Not directly examined in this study, but still worth mentioning, is the importance of the position of additional assistant personnel in the room. Their position should be outside the maximum aerosol trajectory. A effective technique to further reduce the exposure, laminar flow top-to-bottom ventilation could be used which is already often installed in surgical rooms.

## Acknowledgements

The authors acknowledge support from the Central Institute for Scientific Computing (ZISC) and computational resources and support provided by the Erlangen Regional Computing Center (RRZE). The projected was supported by Bayerisches Wissenschaftsministeriums.

